# Application of fuzzy measures to move towards the cyber-taxonomy

**DOI:** 10.1101/2023.03.06.531255

**Authors:** Richardson Ciguene, Julien Mozziconacci, Francis Clément, Aurélien Miralles

## Abstract

The species inventory of global biodiversity is constantly revised and refined by taxonomic research, through the addition of newly discovered species. This almost three century old project provide a knowledge foundation essential for humankind, and notably to develop appropriate conservation strategies. This task relies on the study of millions of specimens housed all around the world in natural history collections. Since two decades, taxonomy generates a plethoric amount of numeric data every year, and notably through the digitization of collection specimens, gradually transforms into a big data science. In this line, the French National Museum of Natural History (MNHN) has embarked into a major research and engineering challenge within its information system, in order to facilitate the transition towards cyber-taxonomic practices which require a facilitated access to data on reference collection specimens housed all over the world. To this end, a first mandatory step is to automatically complete classification data usually associated to collection specimens found in multiple databases. We use here fuzzy approaches to connect one database with other databases and match identical specimens found in each databases together.

## I. Introduction

Species are regarded as the fundamental units for many initiatives related to biodiversity (eg. research, legislation, conservation, agriculture, health...). Around 2 million species have been discovered so far, and 15,000 to 20,000 new species are described and named every year by taxonomists [1] [2]. Nevertheless, only 1 to 10% of the “real” diversity would have been described so far. At the current rate, it would take many centuries to complete the inventory of life diversity. Faced with the magnitude of the upcoming biodiversity crisis, taxonomic research must urgently engage in an “industrial transition” of its research practices.

The herculean task that represents the inventory of global biodiversity fundamentally relies on the study of large international reference collection, which currently house more than 3 billion specimens around the world [3]. With today’s methodological innovations, scientists are increasingly generating diversified dataset necessary to characterize and compare the specimens they are studying (non exhaustively, 2D or 3D images, -omics data, specimen metadata or publications, see also figure 1)).

**Fig. 1.**
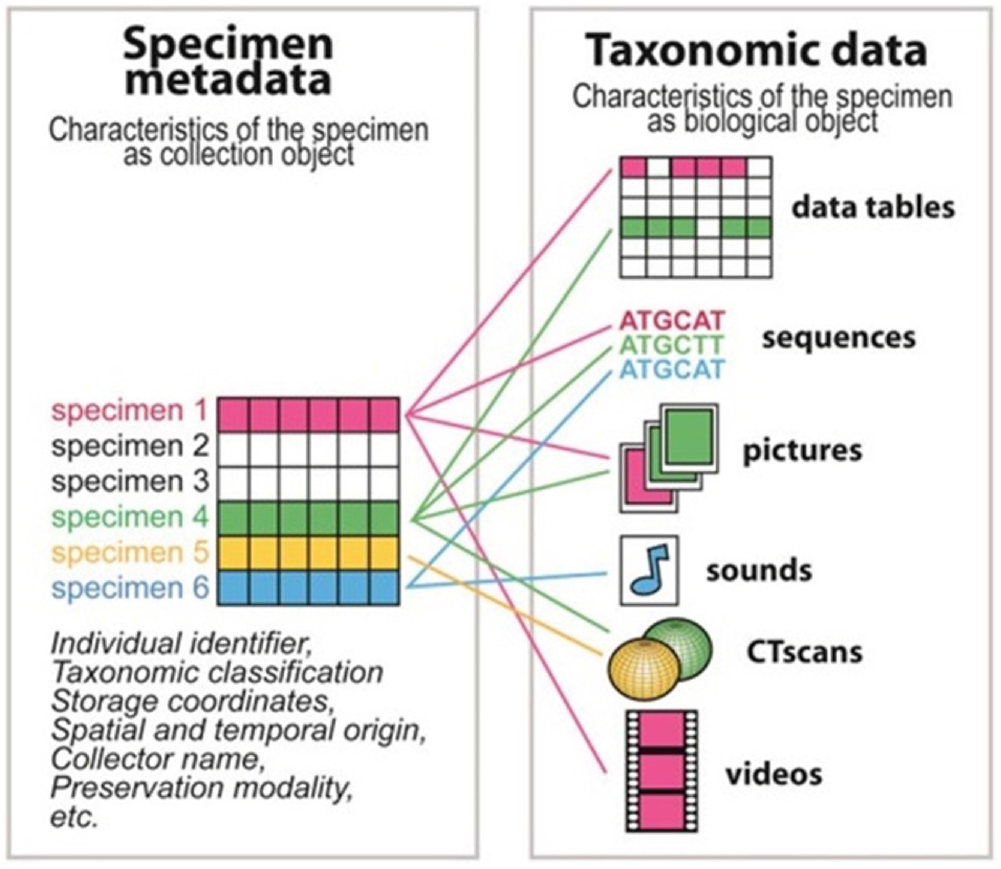
Variety of data composing “cyber-specimens” [8].

Based on the volume of data generated, their variety and their velocity, taxonomy can today be qualified as a big data science [4]. Although a considerable amount of data have been generated and deposited in various repositories, little efforts have yet been made to better synchronize them and make these data FAIR (Findability, Accessibility, Interoperability, and Reusability) [5] [6]. Consequently, the notion of cyber-specimens represents today more an ideal to be reached than a really exploitable resource from a taxonomist perspective.

### A. The linnean classification: a need for fuzzy methods

The linnean classification [20] of organisms consists in a series of hierarchically nested assignments of a specimen to taxon of higher ranks (eg. Genus, Family, Order) (Figure 2).

**Fig. 2.**
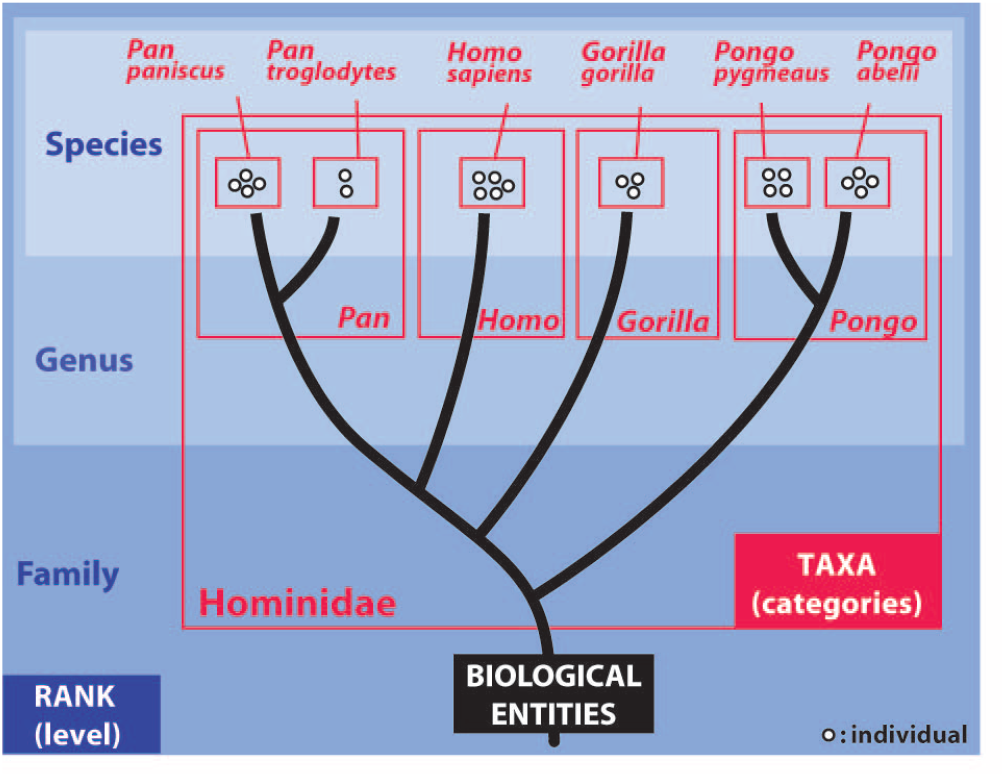
Illustration of the classification of great apes. Classification data in red, with the corresponding linnean ranks in blue. [7].

These multiple assignments constitutes the classification data associated to a specimen. This information is essential for biodiversity experts, as it guides them through the maze of the world’s biodiversity, composed of millions of described species. Unfortunately, the quality of classification data in databases is extremely variable (e.g. errors, inconsistencies or empty data). These shortcomings may severely limit researchers’ ability to harvest a maximum of the data available in various repositories, and impede the future development of fully integrative cyber-taxonomic investigations.

To date, there are few applications of fuzzy methods in such a context. Some taxonomic aggregators like GBIF use fuzzy measures to match taxonomic data from various sources. Also, it allows access to data from their database without necessarily knowing the identifiers of the specimens, but by submitting information such as its scientific name [21]. Basically, these algorithms make use of fuzzy methods, in a data quality context, but most for internal use.

Another example of work like Taxamatch [22], which exists since 2008, proposes a taxa matching method based on phonetic similarity approaches. However, it does not necessarily fit into the context of the quality of classification data as we propose here. On the other hand, we could use it as a point of comparison to assess the accuracy of our work.

Our work aims to experiment with the application of fuzzy methods in taxonomy in two main ways. On the one hand, to improve the quality of the specimen classification data. On the other hand, to effectively match specimen data and interconnect various data sources around the same specimen, which is the core of cyber-taxonomy. For these specific purposes, it is necessary to use the correct distance metric to assess similarity between specimen data. Since the good interconnection of data sources depends largely on the quality of the classification data, in the rest of the paper, we will only focus on the quality upgrading task.

## II. Methods

By data quality upgrading, we mean strengthening the classification data of specimens in our various naturalist databases, through actions of duplication, cleaning and enrichment.To achieve this goal we wish to cross our data with taxa aggregators or taxa referential as well as to cross our databases with themselves for the detection of duplicates. In both cases, we need a method to measure the distance between the taxa to which the specimens are assigned.

### A. Sate-of-the-art distance measures

Number of distance measures can be used. The Hamming distance [11] [12] which quantifies the differences between two sequences of symbols of the same length as the number of symbols, at the same position, that differs. For example the Hamming distance between “rose” and “ruse” is 1, while the Hamming distance between “110110” and “000101” is 4. The Levenshtein distance [13] [14] which corresponds to the minimum number of characters that must be deleted, inserted or replaced to go from one sequence to another. On the other hand, there are various methodologies from Natural Language Processing (NLP) which allow to carry out textual or phonetic matching [16] [17] [18]. In this work, we leverage NLP libraries which naturally include the different distance measurements are used in the context of this work.

### B. Crossing databases

We consider now two datasets in which each record corresponds to a specimen to which a specific taxa is assigned. The first dataset *A* contains n specimens, and the second dataset *B* contains m specimens. One record of the datasets (e.g. *A_i_* or *B_j_*) contains: its taxonomic ranks (Kingdom, Phylum, Class, Order, Family, Genus and Specific epithet); author information; the vernacular name of the specimen; and additional information (subclass code, citations, location and comments). The question to be answered is: what are the similar, disparate or uncertain taxa assignments between *A* and *B* ?

To answer this question, we compute the Levenshtein distance between each record *A_i_* and *B_j_*. The Levenshtein distance is better suited in this context since it can be used when sequences do not have the same length. Also, it is already implemented in a large number of data science libraries and widely used in multiple research areas.

### C. Similarity score

In practice, taxa associated to one specimen can be longer or shorter, or can be found in different order, even when the two specimens are identical. To account for these possibilities, we first aggregate the different fields of interest into on string of characters. We then reorder these characters according to an arbitrary alpha numerical order. We then compute the pairwise Levenstein distance *Dl_ij_* between *A_i_* and *B_j_*. We finally compute a similarity score between 0 and 100 corresponding to (100-*Dl_ij_*)/*L_min_* where *L_min_* is the length of the smallest string of characters.

## III. Practical case and results

### A. A case study: the MNHN databases

To assess the global data challenge on specimen classification data, it is important to remember that the National Museum of Natural History hosts specimen data collected of more than 200 years and that some biodiversity aggregators like GBIF (Global Biodiversity Information Facility) host over 2 billion specimens. In order to clean and aggregate this disparate data into a unified datahub [9] [10] we first need to homogenize the databases of naturalist specimen collections. We illustrate here the first steps of this project, using the database of “Reptiles and Amphibians” collection specimens which contains the following data tables:

- Specimen: contains the basic information of the specimen.
- Origin: contains information on its origin, its harvest or its collection.
- storage: contains information about its storage.
- Loan: who manages their loans for different reasons such as exhibitions and studies.
- Determination: contains information related to its taxonomic description and classification.

In order to move towards cyber-taxonomy and to relate the specimens found in this database with specimens found in other databases, we focus on the “Determination” table, which contains the classification data, a.k.a taxa. This table, which includes “specific epithet”, “genus” and “family” fields is the most useful to establish links between the different data sources. Two important difficulties need to be overcome to match taxa from specimens found in different databases together. Firstly, the assignments to the taxa are not always complete and some fields (or “ranks”) can sometimes even be empty. secondly, these fields can also contain spelling errors. To overcome these issues, fuzzy matching methods appear as a method of choice and we will present hereafter an application.

### B. Experimental protocol

To conduct this experiment, we compared, on the one hand, the “Reptiles and Amphibians” dataset that contains 12811 records and, on the other hand, the Global Biodiversity Information Facility (GBIF) aggregator database [19] which contains about 7 million records, with the same database schema.

Technically, to make the comparisons, we do not use all the variables of the tables. Based on the recommendations of taxonomist experts and our own experience we decided to use only the fields “Genus”, “specific Epithet”, and “scientific Name, i.e. full linnean binomen”. Indeed, this information is sufficient to know if the assignments of taxa are similar or not for two specimens. In figure 4 we provide examples that illustrate the fields extracted and used for specimen matching from the two datasets.

**Fig. 3.**
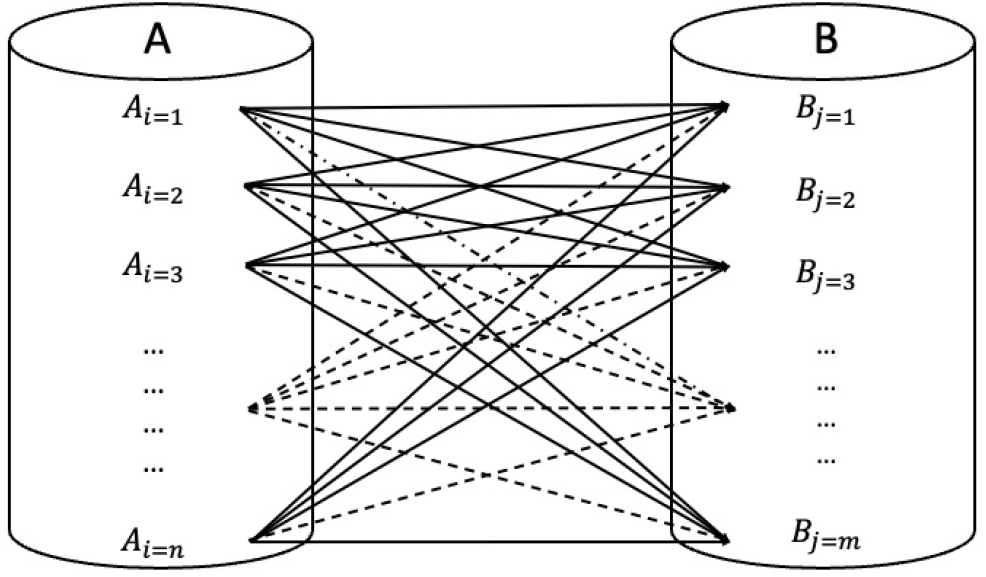
Illustration of our crossing schema between two databases: each record *A_i_* is compared with each record *B_j_*

**Fig. 4.**
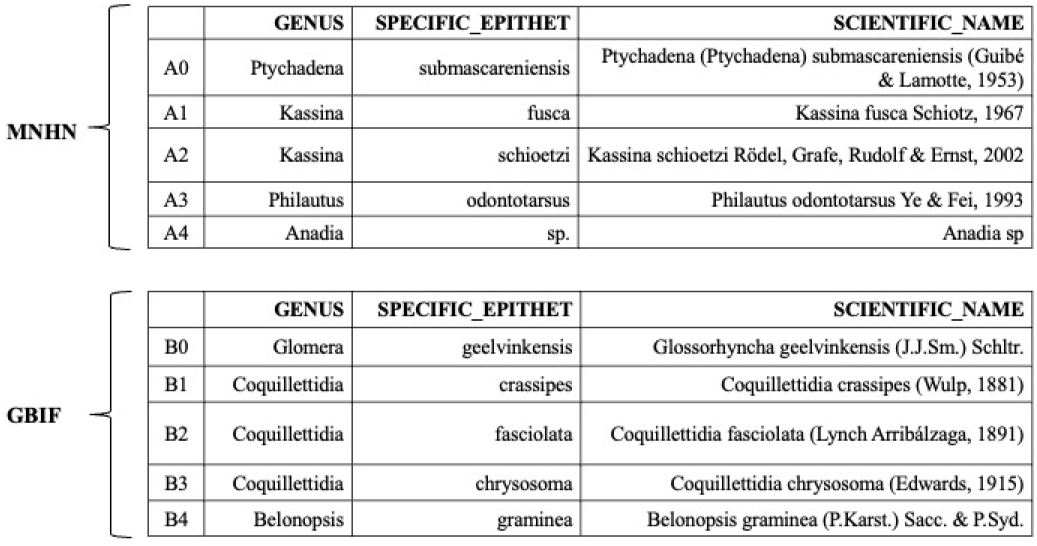
Subsets of the two datasets (MNHN and GBIF) to cross.

### C. Identifying matching pairs of specimens

The highest similarity score for each of the 12811 records was retained for further analysis. The distribution (Fig. 5a) highlights the high number (3242) of perfect matches (similarity scores of 100, in green). However, for most of specimens from the MNHN database (8448, in red) no match was found. For a limited number (1120, in grey), the matching is uncertain: the fields under consideration are not identical but this could be due to spelling errors or to some missing information. We systematically inspected pairs for which the best scores lies between 94 and 99 since the pairs of records exhibiting scores lower than 94 where always characterised as different or potentially misleading according to a taxonomy expert.

**Fig. 5.**
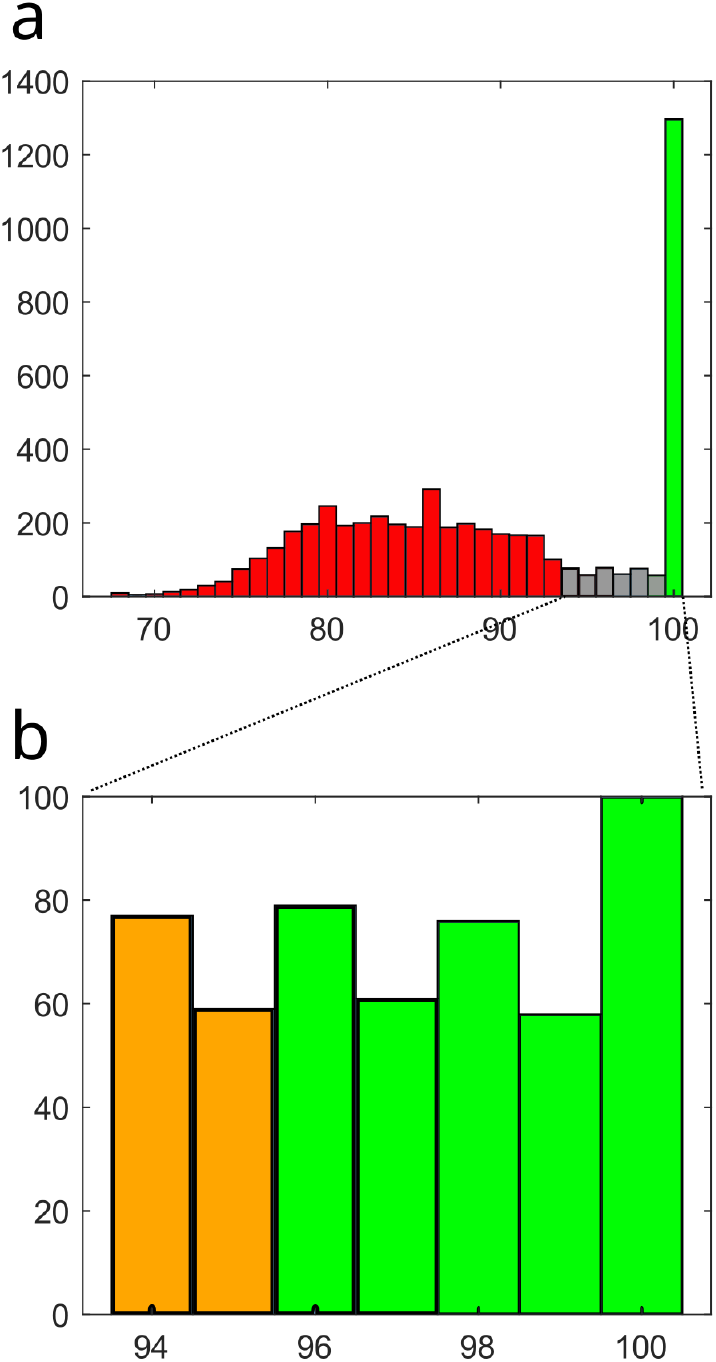
final classification and zoom on incertain area distribution.

We then zoomed in on the grey zone in order to better characterise the small discrepancies that can lead to scores higher than 94. (figure 5.b). Indeed, these scores correspond to three types of differences:

- interval [98, 99]: the typographical errors such as an extra “space” a missing ‘dash”.
- interval [96, 97]: we begin to have more important differences such as errors on the dates which appear in the scientific names or different declension of specific epithets.
- interval [94, 95]: the differences are more pronounced, such as the absence of the author’s middle name in the scientific name, or a different genus name (potentially a synonym).

Analysis by experts in the field led to the classification of the pairs with a score above 95 as similar (in green on figure 5.b) while the pairs with a score of 94 and 95 were uncertain and require human examinations (in orange on figure 5.b). Our procedure thus reduces the human arbitration work required to only 2.7% of the size from the base and further work would be needed to reduce this number.

### D. Follow-up: enriching databases

In figure 6, it is shown a result extract for three MNHN records and their best matches in GBIF. MNHN data is on the left side and GBIF data is on the right side. It should be noted that we could enrich a lot of empty fields on the MNHN side, including the upper taxonomic ranks (i.e. fields highlighted in green in GBIF to fields highlighted in red in MNHN).

**Fig. 6.**
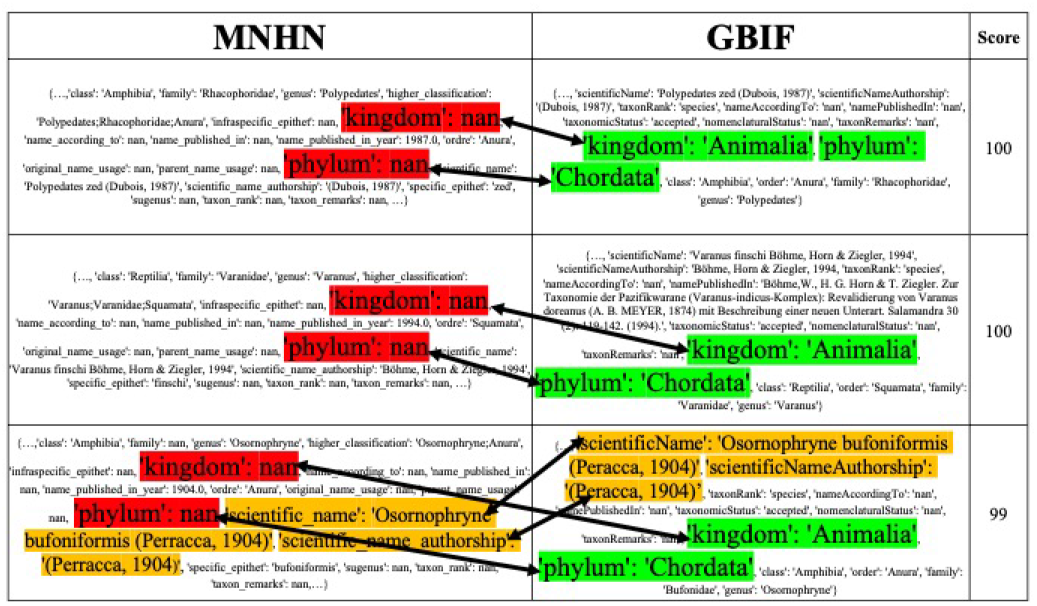
example variables and data from GBIF dataset.

For the third line, with 99% similarity, the difference is in the “Scientific Name”, in the author name with “peRRacca” in MNHN vs “peRacca” in GBIF. So, this could possibly be considered a mistake on MNHN’s side, but that differentiation isn’t so great to say that the two aren’t similar. Thus, the fields highlighted in orange in the MNHN can be replaced/corrected by the fields highlighted in orange in GBIF.

## IV. Conclusion and perspectives

In this article, we have investigated how fuzzy algorithms can participate to the global goal of crossing and unifying naturalist databases in order to move the field towards cyber-taxonomy.

It posed the first formalization of our approach of similarity measure between classification data of specimens to improve their reliability and their level of informativeness, thereby facilitating access to taxonomic data. Beside this preliminary experimentation, many perspectives are planned to test and refine our methods, either in order to extend it to term on all the collections of the MNHN (eg. invertebrates, plants, fungi, microbes), or by replicating comparisons analyses with other repositories or aggregators than the GBIF.

In the short term, refinement work will improve the existing algorithm. Also, we will work on the duplication part of the quality upgrading where we will have to cross our databases with themselves. In the longer term, fuzzy matching methods will be used to link several data sources around the specimens.

The application goal of this work is to avoid human intervention on the entire database, but only on the ‘‘uncertain” class.

The results presented are extremely promising, as only 2,7% of the classification data in the test database will finally require a human arbitration work. Nevertheless, this residual area of uncertainty represent a considerable amount of data to correct manually. To this end, we plan in the medium term to benefit from the expertise of our institutional experts in taxonomy to control and correct our databases. These high quality datasets will then serve as training data for a classification model in order to propose pre-validation on the entire uncertain area from the first treatment. Further, in order to help with data entry, a reinforcement learning model can start from the classification model, in order to learn as it goes to better help in the entry of classification data on specimens.

Since our method is conclusive, we do not exclude the possibility of making it available to other natural history museums and all institutions that have databases on biodiversity.

## Supporting information

Similarity-Output

## Glossary

| N° | Description | Name |
| --- | --- | --- |
| 1 | Plural Taxa, group of one or more populations of an organism or organisms seen by taxonomists to form a unit. (e.g. *Homo sapiens*, Reptilia, Plantae) | Taxon |
| 2 | Hierarchical position of a taxon in the classification (ex. rank of *Homo sapiens* = "species", rank of Reptilia = "Order") | Rank |
| 3 | Individual-based (=specimen-based) data used by taxonomists (ex. morphological or molecular data | Taxonomic data |
| 4 | Assignment of a specimen to a taxon (e.g. to a given species such as "*Homo sapiens*", or to a given family such as "*Hominidae*") | Assignment data |
| 5 | Series of hierarchically nested assignments of a specimen (tacking into account the different ranks for which this information is available (e.g. *Mammalia/Primata/Hominidae/Homo/Homo sapiens*) | classification data |
| 6 | Taxonomic reaserch approach mobilizing all the available digital data obtained specifically from the specimen, including photographs of the specimen "in life," morphometric data, genetic (genomics) sequences, sound files, or chemical profiles, all linked by the specimen identifier to the physical/true specimen | Cyber taxonomy |

Thanks to the datahub project team: Lelia N., Philippe L., Franck B., Romeo A., Gary D., Clement O., Quentin W. and Jonathan G.

## Notes

### Competing Interest Statement

The authors have declared no competing interest.

